# SMAP: A pipeline for sample matching in proteogenomics

**DOI:** 10.1101/2021.09.17.460682

**Authors:** Ling Li, Mingming Niu, Alyssa Erickson, Jie Luo, Kincaid Rowbotham, He Huang, Yuxin Li, Yi Jiang, Chunyu Liu, Junmin Peng, Xusheng Wang

**Affiliations:** Department of Biology, University of North Dakota, Grand Forks, ND 58202, USA; Departments of Structural Biology and Developmental Neurobiology, Center for Proteomics and Metabolomics, St. Jude Children’s Research Hospital, Memphis, TN 38105, USA; Central Laboratory of Zhejiang Academy of Agricultural Sciences, Zhejiang Academy of Agricultural Sciences, Hangzhou 310021, China; Department of Epidemiology and Biostatistics, School of Public Health, Tongji Medical College, Huazhong University of Science and Technology, Wuhan 430030, China; Department of Psychiatry, SUNY Upstate Medical University, Syracuse, New York, 13210, USA

**Keywords:** Sample mix-up, verification and calibration, mass spectrometry, proteomics, proteome, omics, high-throughput technologies, proteogenomics

## Abstract

Integration of genomics and proteomics (proteogenomics) offers unprecedented promise for in-depth understanding of human diseases. However, sample mix-up is a pervasive, recurring problem, due to complex sample processing in proteogenomics. Here we present a pipeline for <u>S</u>ample <u>Ma</u>tching in <u>P</u>roteogenomics (SMAP) for verifying sample identity to ensure data integrity. SMAP infers sample-dependent protein-coding variants from quantitative mass spectrometry (MS), and aligns the MS-based proteomic samples with genomic samples by two discriminant scores. Theoretical analysis with simulation data indicates that SMAP is capable of uniquely match proteomic and genomic samples, when ≥20% genotypes of individual samples are available. When SMAP was applied to a large-scale proteomics dataset from 288 biological samples generated by the PsychENCODE BrainGVEX project, we identified and corrected 18.8% (54/288) mismatched samples. The correction was further confirmed by ribosome profiling and assay for transposase-accessible chromatin sequencing data from the same set of samples. Thus our results demonstrate that SMAP is an effective tool for sample verification in a large-scale MS-based proteogenomics study. The source code, manual, and sample data of the SMAP are publicly available at https://github.com/UND-Wanglab/SMAP, and a web-based SMAP can be accessed at https://smap.shinyapps.io/smap/.

## 1. Introduction

With the development of high-throughput technologies in recent years, remarkable accomplishments have been made by well-conceived and large-scale projects carried out by large consortia, such as the Cancer Genome Atlas (TCGA) project ^1^, the Clinical Proteomic Tumor Analysis Consortium (CPTAC) project ^2-5^, the Genotype-Tissue Expression (GTEx) project ^6^, and the Encyclopedia Of DNA Elements (ENCODE) Project ^7^. These studies usually involve collecting single- or multi-layer omics data from a large number of subjects, followed by statistical significance tests. The power and effectiveness of the statistical tests rely on the accuracy of sample identity ^8^. However, sample mix-up in large-scale multi-omics studies, as a result of specimen labeling errors, and data mismanagement, is a widespread problem ^9^. Such error is often neglected, and it generally leads to irreproducible results, weakened statistical testing power, and false conclusions ^10^. Therefore, to reduce the bias and increase the precision for subsequent analyses, verification of sample identity is one of the first steps in a large-scale omics project.

Mass spectrometry (MS)-based proteomics has emerged as an important molecular profiling technology ^11^ that is complementary to other omics, such as genomics and transcriptomics. Due to recent technical advances, MS-based proteomics has undergone rapid development yielding numerous large-scale proteomics data ^2-5^. Most of these data are generated by multiplexed isobaric labeling-based protein quantification methods, such as isobaric tags for relative or absolute quantitation (iTRAQ) and tandem mass tag (TMT). For example, the TMT-based strategy can now measure as many as 27 samples in one batch simultaneously ^12^, and potentially allows for the measurement of 81 samples when combined with metabolic labeling by amino acids in cell culture (SILAC). These multiplexing strategies would further exacerbate the sample mix-up problem, posing a significant challenge for sample verification and calibration.

Several methods have been developed to verify sample identity in large-scale genomics and transcriptomics studies using genotype concordance ^13^ and correlation of variant allele fractions ^14^. We have also implemented a genotype-based method to process sample mix-up problem from sequencing-based transcriptomic data ^15^. Most of these methods exploit the allelic information of a sample, which can be directly derived from sequencing data. Calling genotype from proteomics data from multiplexed isobaric labeling-based quantitative proteomics data remains a major challenge for several reasons because multiple labeled samples are further mixed during the experiment. While the proteogenomics approach has been widely used to detect variant peptides, it only call variant peptides from label-free data or mixed samples ^16-18^. As a result, there is no method available for verifying and calibrating sample identity in a multiplexed quantitative proteomics study.

In this study, we describe a pipeline for <u>S</u>ample <u>Ma</u>tching in <u>P</u>roteogenomics (SMAP). SMAP first performs the proteogenomics database search to detect variant peptides from multiplexed isobaric labeling-based quantitative proteomics data, and then infers allelic information of each sample based on its expression level of the variant peptides. SMAP finally verifies and calibrates sample identity based on a combination of concordance and specificity scores using inferred allelic information. The performance of SMAP is assessed by a simulation study and a large-scale proteomics dataset.

## 2. Results

### 2.1 Method implementation

#### 2.1.1 Identifying variant peptides

SMAP consists of three main components (**Fig. 1** and **Supplementary Figure 1**): (i) identifying variant peptides using the proteogenomics approach; (ii) inferring allelic information for each sample; and (iii) verifying and calibrating sample identity. SMAP can take quantification table of variant peptides from the proteogenomics tool. In this study, we uses JUMPg, to identify variant peptides ^18^. JUMPg constructs a customized database using genomic variant files (e.g., VCF files) or indirectly from RNAseq raw data (FASTAQ file), in which JUMP performs preprocessing, tag generation, MS/MS pattern matching, and scoring as previously reported ^19^. The identified variant peptides are further filtered with the target-decoy strategy to control false discovery rate (FDR) ^20,21^.

**Figure 1.**
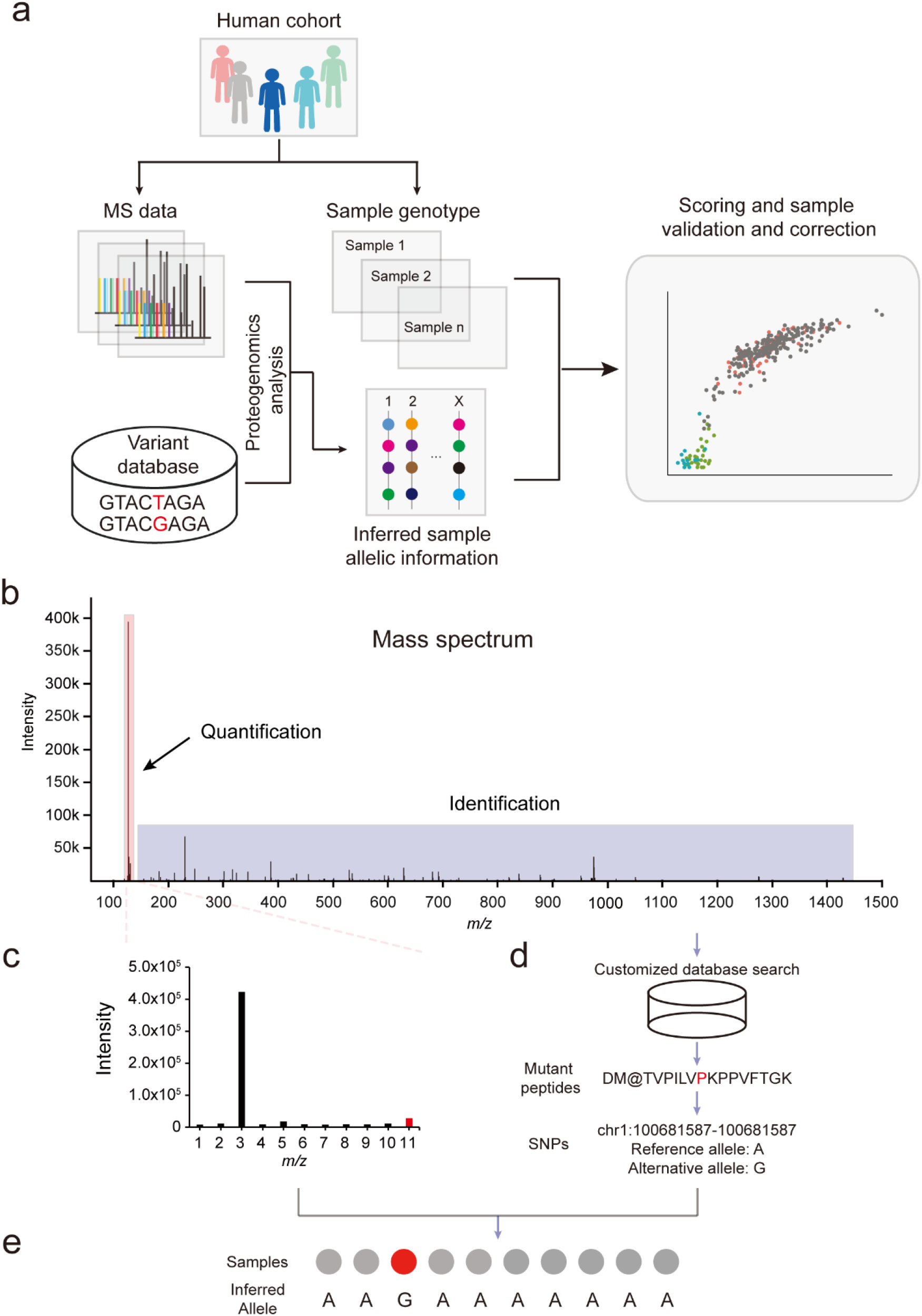
Architecture of SMAP. (a) Schematic diagram of the SMAP pipeline. Mass spectrometry (MS)-based proteomics data are generated from a large-scale project. The data are searched against the customized database containing all theoretical variant peptides generated from whole-genome DNA sequencing, RNA sequencing, or public databases (e.g., dbSNP). For each batch of the TMT-based quantitative proteomics data, variant peptides are identified through the proteoteogenomics analysis. Two scores (i.e., concordance score and delta concordance score) are computed for verifying and calibrating the sample identity. (b) Example of a MS spectrum containing TMT report ions. (c) A zoom-in view of intensity of reporter ions shown in panel A. (d) Identification of variant peptides using the proteogenomics approach. (e) Inferring sample alleles from quantitative expression level of the variant peptide for each sample.

#### 2.1.2 Inferring allelic information for each sample

SMAP infers sample allelic information based on the relative expression level of each sample (**Fig. 1a** and **Supplementary Figure 1**) for variant peptide identified in proteomics data with multiplexed isobaric labeling-based quantification methods. For each identified peptide, the intensity of reporter ions is extracted as the expression level for each sample (**Fig. 1b**). The intensity of each sample is transformed to log_2_-scale, followed by a scale normalization (**Fig. 1c**) using the following formula,

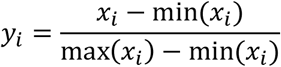

where *x*_*i*_ and *y*_*i*_ are raw and scaled intensity for a sample, respectively. The scaled intensities are in the range of 0 to 1.

SMAP then infers allelic information of each sample based on the scaled intensity of identified variant peptides (**Fig. 1d**). For a diploid organism, such as human or mouse, there are three possible alleles of a variant peptide: homozygous reference allele (AA), homozygous alternative allele (BB), and heterozygous allele (AB). To determine the allelic information of each sample of a variant peptide, SMAP divides the scaled intensities into three quantiles: lower-quartile (LQ, 25^th^ percentile), inter-quantile (IQ, 25^th^ to 75^th^ percentiles), and upper-quartile (UQ, 75^th^ percentile) (**Supplementary Figure 2**). The three quantiles are assumed to correspond to the reference (AA), heterozygous (AB), and alternative (BB) alleles, respectively (**Fig. 1e)**.

#### 2.1.3 Scoring

Once the allelic information of each sample is inferred for all variant peptides, SMAP verifies and calibrates the sample identity using a combination of two scores: concordance score (Cscore) and specificity score (delta concordance score: ΔCscore) (**Fig. 1a** and **Supplementary Figure 1**). The ΔCscore is defined as the difference between the best concordance score and the following concordance score divided by the best score. Thereby, the ΔCscore is a good measure of separating true from false hits. A confident assignment generally has a high Cscore and ΔCscore. To determine whether a sample can be verified, these two scores are combined and required for calibrating its identity.

### 2.2 Performance of SMAP using simulation data

Sample verification and calibration depends largely on reliable allelic information derived from the quantitative proteomics data. To determine how many reliable alleles in a sample are sufficient for SMAP to verify and calibrate its identity, we conducted a simulation study using a subset of genotypic data that were generated by whole-genome sequencing and microarray chip (**Fig. 2a)**. To mimic “real” allelic information derived from a large-scale proteomics dataset, we first exacted a subset of the genotypic matrix composed of 500 SNPs from 420 samples, and then randomly selected a sample. The selected sample was then shuffled with a certain probability (e.g., 10%, 20%, 40%, and 80%). We examined whether the sample identity assigned by SMAP matched the original one based on the Cscore (**Fig. 2a**). As shown in **Figs. 2b-d**, SMAP is capable of successfully validating the sample identity with the reliable genotypic number as low as 20% (i.e., 80% shuffled).

**Figure 2.**
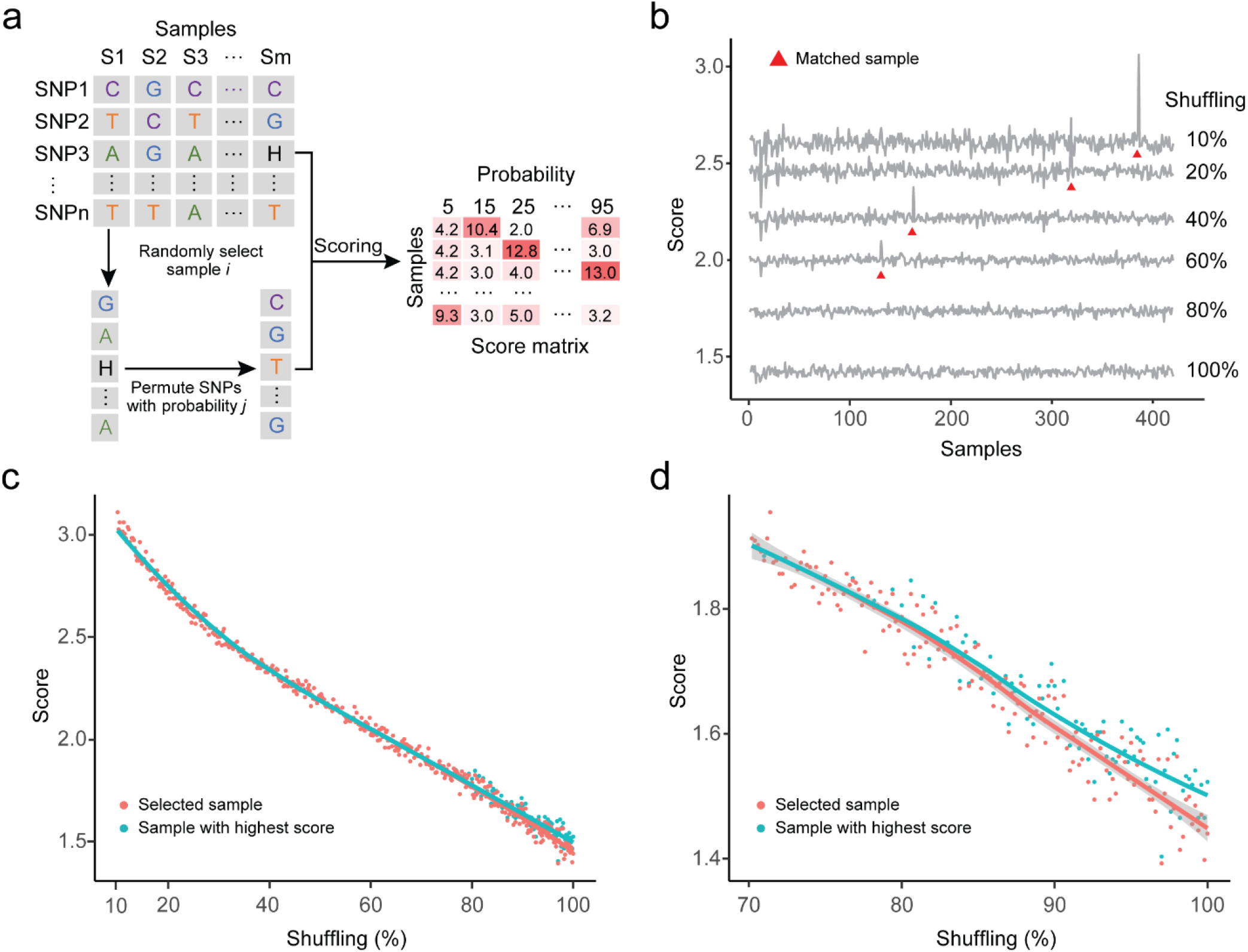
Parameterization and evaluation of SMAP performance using a simulation study. (a) Diagram showing the simulation procedure. The procedure includes 4 steps: (1) A reduced matrix (e.g., 500 rows x 420 columns) is randomly extracted from the reference genotypic data; (2) randomly select a sample; (3) permute the genotypes with a certain probability; (4) A scoring matrix is generated by the SMAP. (b) Curve plot showing the matching score distribution. The X-axis represents 420 samples and the y-axis represents the matching score between a testing sample and each sample. (c) Score distribution of the selected sample with the highest score and the select (i.e., “true”) sample along the percentage of shuffled samples. The X-axis denotes the score and the Y-axis represents the percentage of shuffled samples. The red dot is the selected sample and the blue is the one with highest score. (d) A zoom-in plot of the score distribution with shuffled samples from 70% to 100%.

### 2.3 Application of SMAP to PsychENCODE BrainGVEX proteomics data

Next, we applied SMAP to a deep proteomics dataset generated by the PsychENCODE BrainGVEX project, in which 288 biological samples and 31 internal controls (i.e., a mixture of 288 samples) were quantified by 29 batches of 11-plex TMT-based proteomics technology. In addition to proteomics data, the PsychENCODE BrainGVEX project also generated other omics data with matched samples, including 285 samples with low-depth whole genome sequencing (WGS), 426 samples with RNA sequencing (RNA-Seq), 295 samples with assay for transposase-accessible chromatin using sequencing (ATAC-Seq), and 197 samples with ribosome sequencing (Ribo-Seq). Previous analysis using DRAMS has identified ∼19% mixed samples in both WGS and ATAC-seq, ∼25% in Ribo-seq, and ∼3% in RNA-seq data ^15^.

To identify variant peptides in the PsychENCODE BrainGVEX proteomics data, a total of ∼34 million MS2 spectra from 29 batches of TMT experiments were searched against a customized database. The database contains 20,396 reviewed protein sequences from the UniProt database ^22^ and 17,844,001 theoretical peptides translated from variants generated by WGS and RNA-seq data. SMAP identified a total of 5,065 variant peptides (Supplementary Table1), corresponding to 20,129 SNPs at 1% protein FDR (**Fig. 3a**). The number of variant peptides and SNPs show similar trends across 29 batches. On average, a total of 694 SNPs were identified per batch, ranging from 430 SNPs in batch 23 to 901 SNPs in batch 24. To use reliable SNPs, further filtering of SNPs was completed using the minor allele frequency (MAF) of > 1%, identifying a total of 8,358 SNPs with an average of 288 SNPs per batch (**Fig. 3a**).

**Figure 3.**
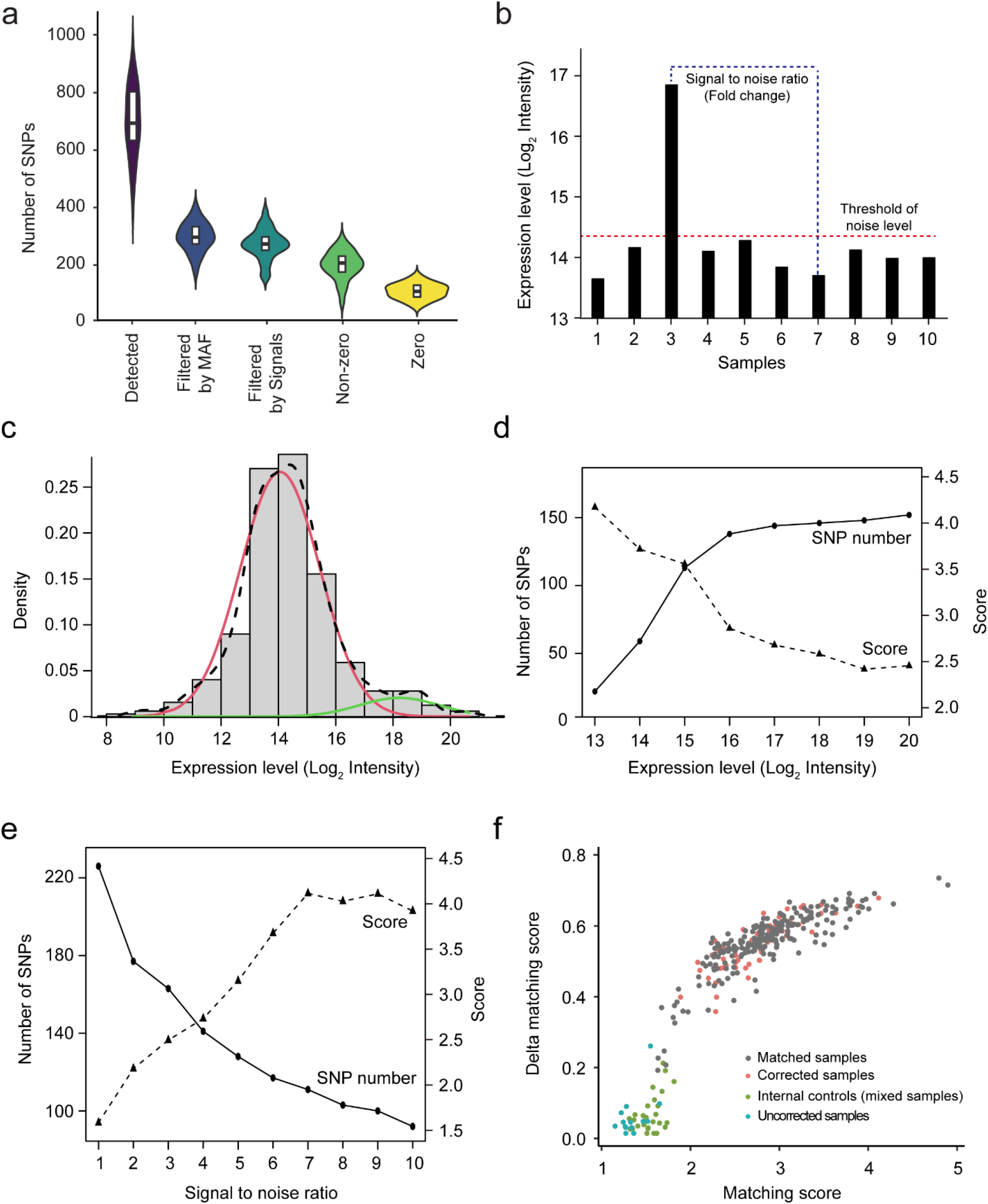
Application of SMAP to the PsychENCODE BrainGVEX proteome data. (a) Number of SNPs identified, filtered by both minor allele frequency (MAF) and threshold of the noise signal, as well as SNPs with and without zero signals detected by the JUMP program. (b) Diagram showing how to define the threshold of the noise background and the signal-to-noise (S/N) ratio. (c) Distribution of minimal signals. Two normal distributions can be separated by the expectation-maximum (EM) method. (d) The distributions of the number of SNPs and the concordance score with different thresholds of the noise background. (e) The distributions of the number of SNPs and the concordance score with different S/N ratios. (d) Score distribution of matched samples, corrected samples, internal controls, and uncorrected samples.

To determine allelic information of a sample for each identified variant peptide, two parameters are critical: a threshold of the noise signal (i.e., intensity of the homozygous reference allele (AA) in the variant peptide), and a signal-to-noise (S/N) ratio (i.e., the signal rnatuatio between the alternative homozygous allele (BB) and the homozygous reference allele (AA)) (**Fig. 3b**). For a variant peptide, the signal of a sample with a homozygous reference allele (AA) is expected to be no different from the background noise because the variant peptide is not detectable in the TMT mixed experiment. To determine the threshold of the background noise level, we used one batch (i.e., Batch 1) of the 29 TMT experiments to determine the distribution of the background noise level of all identified variant peptides. We found that the distribution can be clearly separated into two modes by the expectation-maximum (EM) method, consisting of an authentic noise distribution (mean = 14.06, SD = 1.39; Log_2_ intensity) and a signal distribution (mean = 18.22, SD = 1.29; Log_2_ intensity) (**Fig. 3c**). The result indicates that a log_2_ intensity of 16.14 (noise mean + 1.5 SD or signal mean – 1.5 SD) can be used as a cutoff. This is consistent with the result found when evaluating the impact of the minimal intensity on the number of SNP and the concordance score (**Fig. 3d**). The analysis of the S/N ratio between the alternative homozygous allele (BB) and the homozygous reference allele (AA) showed that an S/N value of 3 achieves a reasonably good number of SNPs and concordance score (**Fig. 3e**). Following filtration by the minimum signal and the S/N ratio, we detected 7,628 SNPs, with an average of 263 SNPs per batch (**Fig. 3a**).

After optimizing the parameters, we used 7,628 filtered SNPs to infer allelic information for 288 biological samples and 31 controls in the PsychENCODE BrainGVEX proteomics data. The allelic information of each sample was then inferred based on its expression level in a batch using the strategy of three quantiles (**Supplementary Figure 2; see method**). To verify the sample identity, we first evaluated the distribution of Cscore and ΔCscore of the 31 internal controls, which were mixed samples that functioned as a negative control. The internal controls showed an average Cscore of 1.50 and a standard deviation of 0.13 (**Supplementary Figure 3a**), and an average ΔCscore of 0.05 and a standard deviation of 0.05. When applying SMAP to the first batch of the proteomics data, the samples can be clearly separated from internal controls and matched samples with the Cscore of 1.50 and ΔCscore of 0.20, identifying one sample as being mixed up (**Supplementary Figure 3b**). For 288 biological samples, SMAP identified 276 biological samples with the Cscore above 1.50 and ΔCscore above 0.20. When comparing those 276 samples with their original labeling identities from DNA-based genotyping data, we found that 54 samples (18.75%) showing mixed-up identification (**Fig. 3f**). In addition, a total of 12 samples showed a score below the threshold, suggesting that these samples could not be calibrated for their identities.

### 2.3 Cross-validation of sample correction

The multi-omics data generated by the PsychENCODE BrainGVEX project enable cross-validation of sample calibration made by SMAP. As the identity of samples has previously been verified and calibrated in other omics data, such as samples used for ATAC-seq and Ribo-seq data ^15^, we next evaluated whether the sample identities in the proteomics data calibrated by SMAP are consistent with that made in both ATAC-seq and Ribo-seq data. We found that 34 out of 54 samples (63%) showed the same calibration in samples from all three platforms: MS-based proteomics, ATAC-seq, and Ribo-seq. In addition, a total of 9 samples with the same calibration between proteomics and ATAC-seq, 5 between proteomics and Ribo-seq, and 6 proteomic-specific calibrations (**Fig. 4a**).

**Figure 4.**
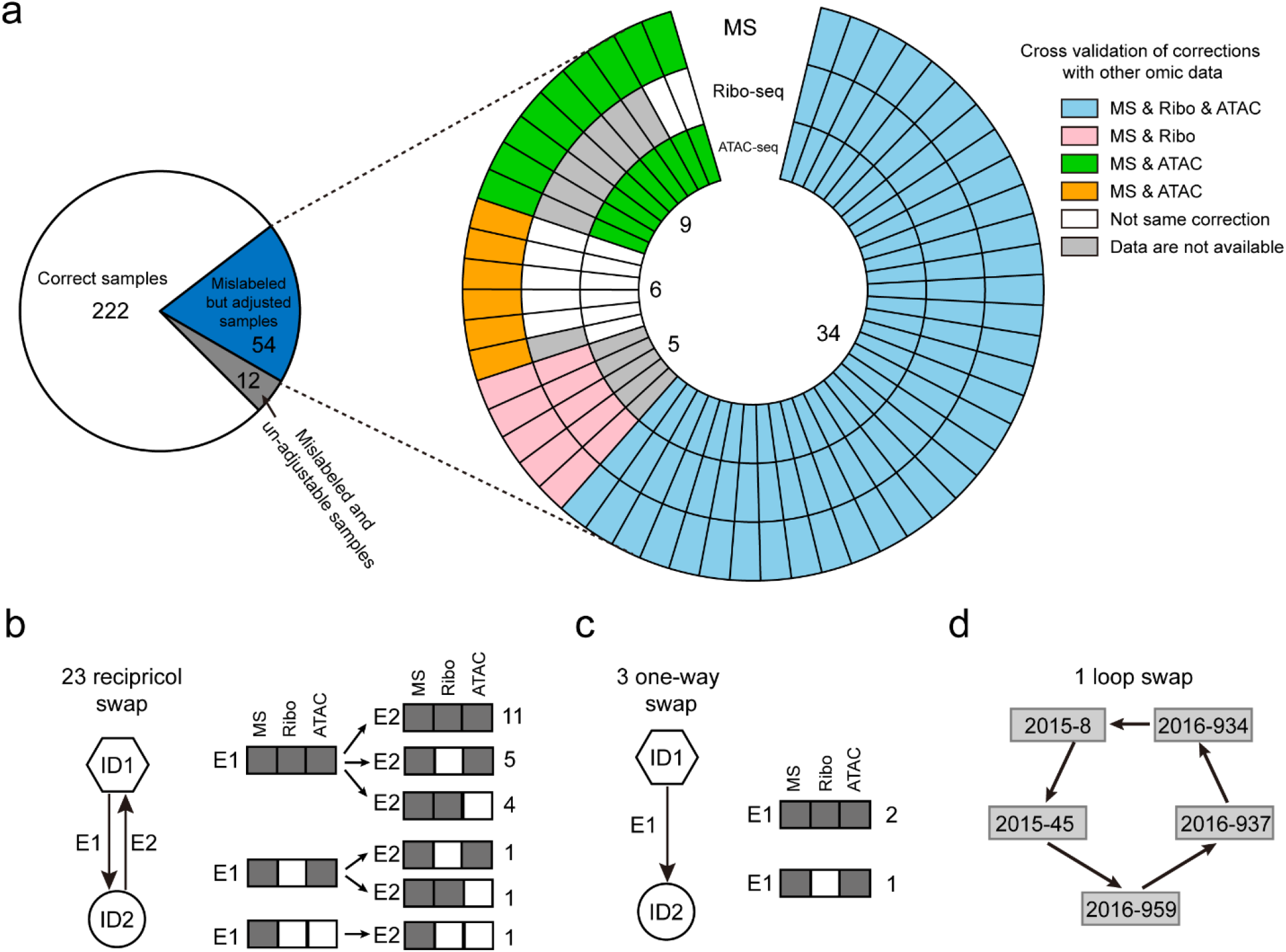
Validation of sample calibration across other omics platforms. (a) The pie diagram shows the number of correct samples, mislabeled but adjustable samples, and mislabeled and un-adjustable samples. The circular plot shows validation of sample corrections across different platforms. Each circle represents an omics platform. (b-d) Three types of sample calibrations and comparison of calibrations across different omics platforms.

We next sought to investigate how these 54 samples were mixed in the proteomics experiment. We summarized the calibrations into three categories: reciprocal, one-way, and cyclic swaps. We found that 46 samples (23 pairs) were reciprocal swapped (**Fig. 4b**), with about half of the samples (11 pairs) supported by the same calibration in all three platforms: MS-based proteomics, ATAC-seq, and Ribo-seq, followed by 9 pairs that were supported by three platforms in one direction and two platforms in the other direction. For the three one-way swap, two calibrations are supported by three platforms and one by two platforms (**Fig. 4c**). Most of the cyclic swaps (5/6) are proteomic-specific calibrations (**Fig. 4d**). The result suggests that sample mix-up is a not random process but follows a certain pattern in the lab procedures.

## 3. Discussion

Sample mix-up is a pervasive problem in almost all large-scale omics studies. In this study, we described a novel method, SMAP, for validating and correcting samples that are used for a large-scale TMT-based quantitative proteomics study. By applying SMAP to proteomics data from 288 biological samples in the pyschENcode BrainGVEX project, we verified and calibrated ∼18% mixed samples. Based on the simulation study, SMAP is capable of validating the sample identity of as few as 20% of the reliable genotypes.

While the proteogenomics approach is increasingly used to detect protein-coding variants and alternative splicing sites ^16-18^, there is no study to infer allelic information for each sample in multiplexed isobaric quantitative proteomics data. It is the first time that SMAP determines allelic information of each sample based on the expression level of the variant peptides. In this study, we also proposed a novel combination score for evaluating sample verification and calibration. With advances in next-generation sequencing and high-resolution mass spectrometry technologies, omics data from the same set of biological samples are now routinely collected in a project. Appropriate data quality controls at each step are required to ensure high data integrity. Therefore, SMAP complements existing approaches for sample verification and calibration that have been developed for genomics and transcriptomics data.

Although SMAP can infer sample-level allelic information for sample verification, the inference certainty was still imperfect. First, the expression level of a variant peptides is quantitative, unlike qualitative allele from genomics and transcriptomics data generated by sequencing technologies. Although we proposed to use three quantiles to determine the allele types, the allelic information inferred by SMAP shows uncertainties. In the analysis of the PsychENCODE BrainGVEX proteomics data, we found many mismatched alleles are in the boundary of each range. Secondly, the homozygous alternative allele (BB) and heterozygous allele (AB) cannot be distinguished solely depending on the expression level if only AB alleles are present in the samples. To mitigate this issue, SMAP examines the number of allele types in the original genotypic data. Thirdly, expression imbalance, such as single-parental expression (e.g., imprinted expression) and over- or under-dominant expression of the identified variant peptides would potentially influence the inference of allelic information ^23^. The level of mix-up detected in our proteomics data is similar to that in other next-generation sequencing-based omics data. Large-scale omics data profiling and analysis typically involves many research laboratories and mistakes could occur in different ways at the significant level. Sample mix-up is not only a common problem but also a major problem getting less attention than it deserves. By comparing data of multi-omics data, we had the best chance of correct sample identifies, which is essential for performing high-quality research.

In summary, we present a robust and easy-to-use method and tool, SMAP, for sample verification and calibration. SMAP can successfully calibrate ∼18% mixed sample in a large-scale proteomics data. The concordance score and delta concordance score are approved to be effective in sample verification. We recommend sample verification and calibration as an important part of the data analysis in a large-scale MS-based proteomics study.

## 4. Methods and Materials

### 4.1 Proteomics and other omics datasets

A total of 288 well-characterized postmortem human brain samples (179 males, 109 females) from the Stanley Medical Research Institute and Banner Sun Health Research Institute were used for this study ^24^. These samples were collected from 211 neurotypical controls, 48 individuals with schizophrenia (SCZ), and 29 individuals with bipolar disorder (BD). The samples include 282 Caucasians, 1 Hispanic, 1 African American, 3 Asian American, and 1 unknown. The brain frontal cortex samples were subjected to 29 batches of deep proteome profiling using TMT LC/LC-MS/MS technology.

In addition to the proteomics data that we generated, five other omics data are also available at the psychENCODE knowledge portal (https://psychencode.synapse.org/Explore/Studies/DetailsPage?studyName=BrainGVEX), including RNA-Seq data from 426 samples (274 males and 152 females), assay for transposase-accessible chromatin using sequencing (ATAC-Seq) data from 295 samples (180 males, 112 females, and 3 unknown-sex samples), and ribosome sequencing (Ribo-Seq) data from 197 samples (122 males, 70 females, and 5 unknown-sex samples). The genotypes were generated by the combination of three platforms, including Affymetrix chip data, whole-genome sequencing, and RNA-seq data ^15^.

### 4.2 Identification of variant peptides in the proteomics data

JUMPg program was used to identify variant peptides by taking MS-based proteomic data and a customized database curated with genomic variant data ^18^. In the JUMPg program, the variant peptides were identified by the JUMP search engine, which performs preprocessing, tag generation, MS/MS pattern matching, and scoring as previously reported ^19^. JUMP was used to search MS/MS raw data against a composite target/decoy database (Junmin Peng, 2003) to evaluate false discovery rate (FDR). The target database contains both core protein sequences downloaded from the UniProt ^22^ and all theoretical variant peptides from all nonsynonymous protein-coding variants. To generate theoretical variant peptides, all variants in the genotypes were re-annotated using the genome annotation tool ANNOVAR ^25^ based on the human reference genome (i.e., GRCh38/hg38 assembly). The decoy database was generated by reversing protein sequences in the target database. Major parameters included precursor and product ion mass tolerance (± 8 ppm), full trypticity, static mass shift for the TMT tags (+ 229.16293) and carbamidomethyl modification of 57.02146 on cysteine, dynamic mass shift for Met oxidation (+15.99491), maximal missed cleavage (*n* = 2), and maximal modification sites (*n* = 3). Putative PSMs were filtered by mass accuracy and then grouped by precursor ion charge state and filtered by JUMP-based matching scores (i.e., Jscore and ΔJn) to reduce protein FDR < 1%. If one peptide could be generated from multiple homologous proteins, the peptide was assigned to the protein with the highest PSM based on the rule of parsimony.

### 4.3 TMT-based Peptide/Protein Quantification by JUMP Software Suite

The analysis was performed in the following steps, as previously reported with modifications ^26^: (i) extracting TMT reporter ion intensities of each PSM; (ii) correcting the raw intensities based on isotopic distribution of each labeling reagent (e.g. TMT126 generates 91.8%, 7.9% and 0.3% of 126, 127, 128 m/z ions, respectively); (iii) removing sample loading bias by normalization with the trimmed median intensity of all PSMs; (iv) calculating the mean-centered intensities across samples (e.g. relative intensities between each sample and the mean), (v) summarizing relative intensities of proteins or variant peptides by averaging related PSMs. The quantification values of variant peptides with and without missing values were extracted for inferring allelic information for each sample by SMAP.

### 4.4 Simulation data

We generated a simulated data to test the performance of SMAP on evaluating how many SNPs identified from MS-based proteomics data are enough for validating and correcting the sample identity. We first randomly extract a subset of genotypic matrix with 500 SNPs and 420 samples. We then randomly selected a sample from the matrix and randomly permutated the allelic information with replacement sampling at a certain probability level *α*. The *α* level was set in the range from 10 to 100 at a step of 1.

### 4.5 Data availability

All raw mass spectrometry files, peptide and protein identification results, and genotypic data are available at the Synapse website (https://psychencode.synapse.org/Explore/Studies/DetailsPage?studyName=BrainGVEX).

### 4.6 Code availability

The SMAP is implemented in both standalone (https://github.com/UND-Wanglab/SMAP) and web-based (https://smap.shinyapps.io/smap/) versions. The standalone SMAP is written in PERL programming language. The web-based SMAP is developed using Shiny, an R package that supports the development of web-based R applications that can be hosted online. Manual, tutorial, and sample datasets are also available at the SMAP project homepage https://sites.google.com/view/smapwanglab/home.

## Acknowledgements

This work was partially supported by two NIH subawards (X.W.): (RF1AG057181), and NIH subaward (1P30DA044223-01). It is also supported by the UND Center for Biomedical Research Excellence (CoBRE) for Epigenomics of Development and Disease (X.W.), the UND CoBRE for Host-Pathogen Interactions (HPI) (X.W.), the ND EPSCoR STEM program (X.W.), and the UND Vice President for Research & Economic Development (VPRED) seed program (X.W.). The MS analysis was performed in the Center of Proteomics and Metabolomics at St. Jude Children’s Research Hospital, partially supported by NIH Cancer Center Support Grant (P30CA021765). NIH R01AG053987 (J.P.), U01MH103340-01, 1U01MH116489, 1R01MH110920 (C.L).

## Author Contributions

X.W. and J.P, contributed to the conception and design of the project. X.W., L.L., J.L., H.H., A.E., and Y.L. developed computational algorithms and wrote the SMAP program. M.N. performed proteomics experiments. C.L. and Y.J. provided human specimens and genotypic data. X.W., L.L., and J.P., wrote the manuscript.

## Supplementary Materials for

### Supplemental Figures and Tables

**Supplementary Figure 1.**
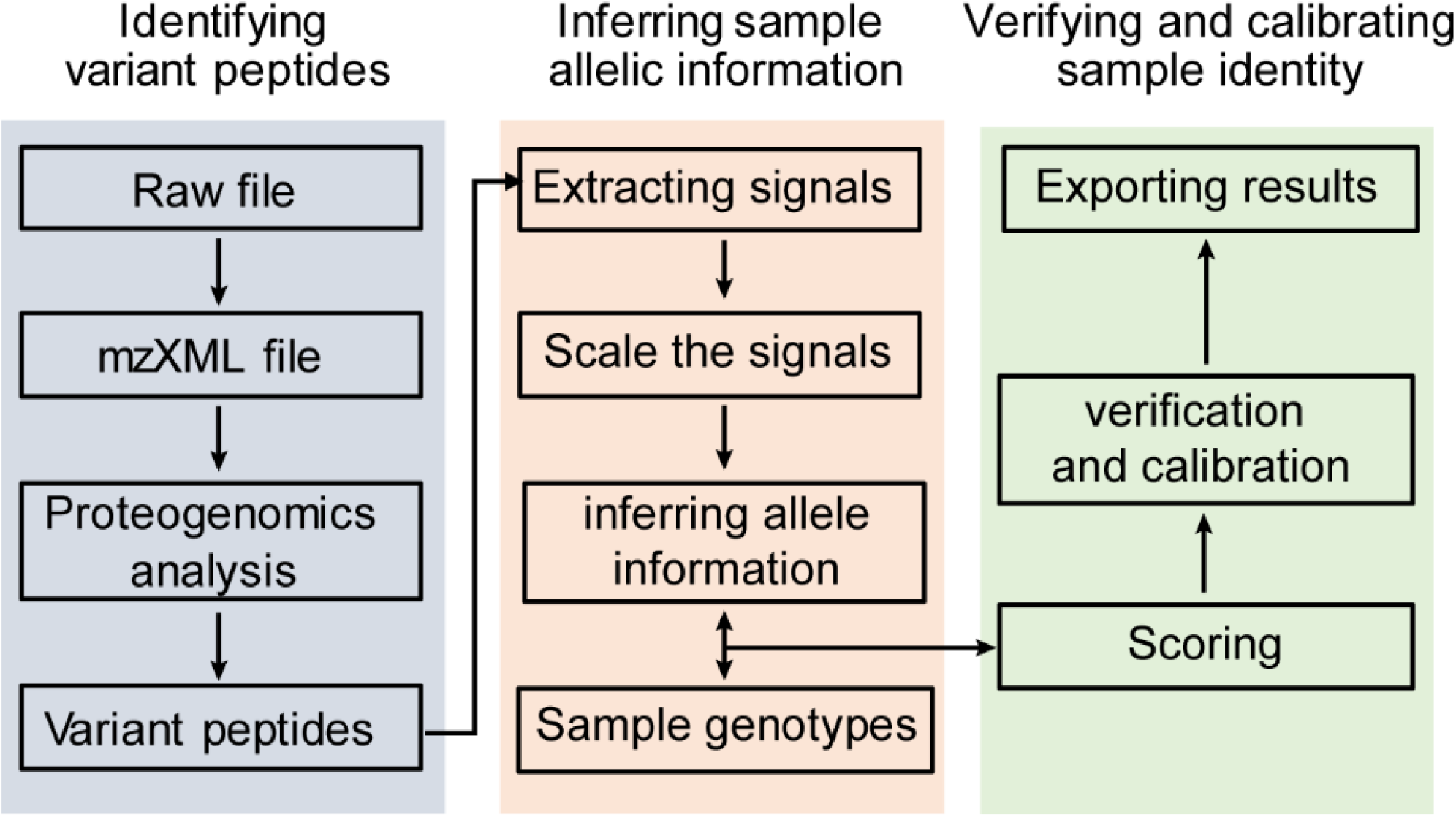
Workflow of the SMAP pipeline.

**Supplementary Figure 2.**
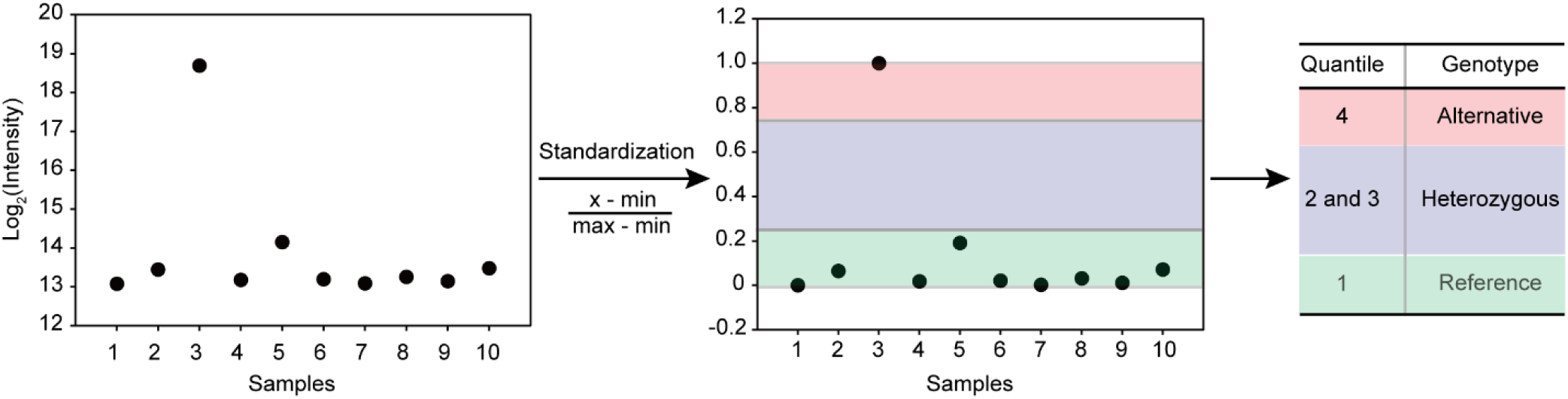
Standardization of intensities and Inferring sample genotypes. The expression levels of a variant peptide was scaled to the range of 0 to 1. The scaled expression levels are quantized into three categories: homozygous reference allele, heterozygous allele, and homozygous alternative allele.

**Supplementary Figure 3.**
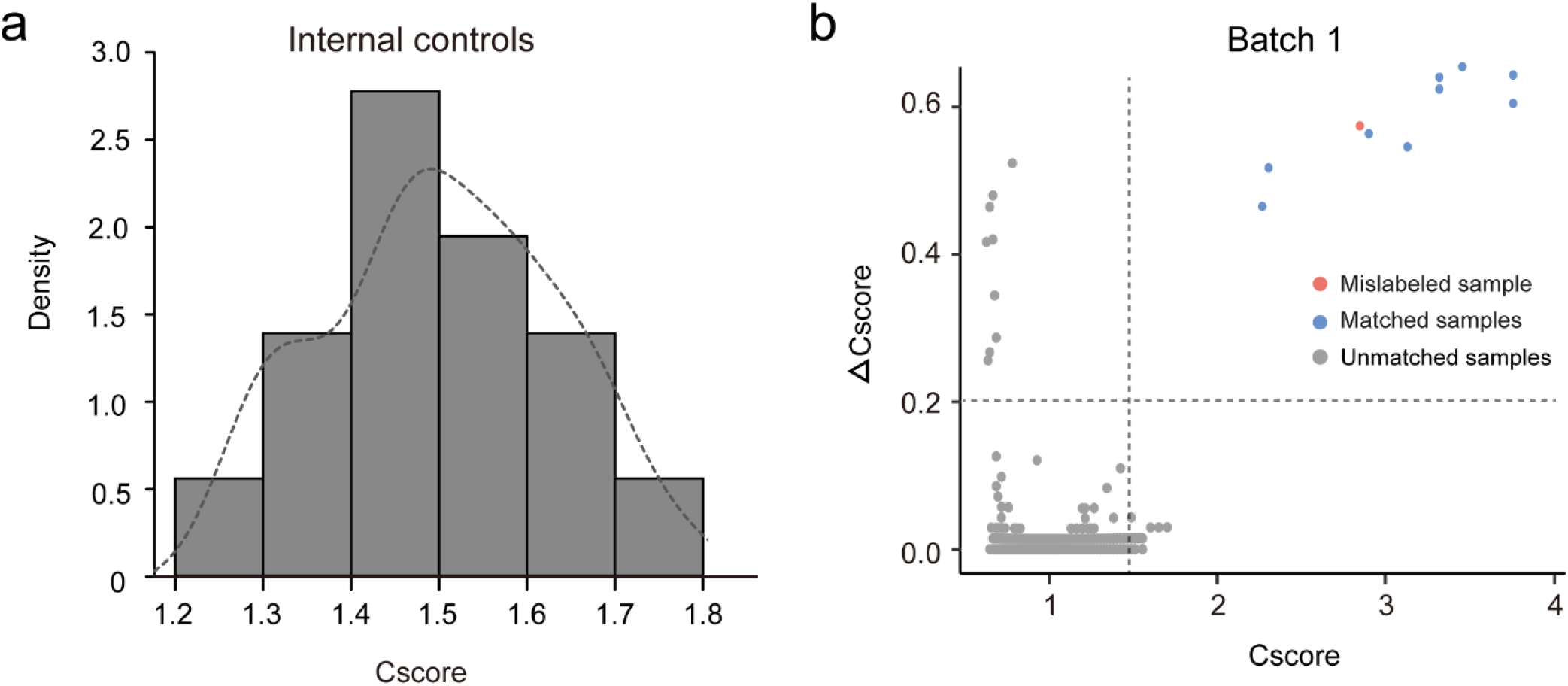
Evaluate of the SMAP pipeline. (a) Distribution of Cscore in internal controls. (b) Distribution of Cscore and delta Cscore when applying SMAP to one batch TMT proteomics dataset.

Supplementary Table 1. One supplemental table in an attached Excel file.

